# Computational approach to dendritic spine taxonomy and shape transition analysis

**DOI:** 10.1101/051227

**Authors:** Tomasz Kusmierczyk, Michal Lukasik, Marta Magnowska, Matylda Roszkowska, Grzegorz Bokota, Dariusz Plewczynski

## Abstract

The common approach in morphological analysis of dendritic spines is to categorize spines into subpopulations based on whether they are stubby, mushroom, thin, or filopodia. Corresponding cellular models of synaptic plasticity, long-term potentiation, and long-term depression associate synaptic strength with either spine enlargement or spine shrinkage. Although a variety of automatic spine segmentation and feature extraction methods were developed recently, no approaches allowing for an automatic and unbiased distinction between dendritic spine subpopulations and detailed computational models of spine behavior exist.

We propose an automatic and statistically based method for the unsupervised construction of spine shape taxonomy based on arbitrary features. The taxonomy is then utilized in the newly introduced computational model of behavior, which relies on transitions between shapes. Models of different populations are compared using supplied bootstrap-based statistical tests.

We compared two populations of spines at two time points. The first population was stimulated with long-term potentiation, and the other in the resting state was used as a control. The comparison of shape transition characteristics allowed us to identify differences between population behaviors. Although some extreme changes were observed in the stimulated population, statistically significant differences were found only when whole models were compared. Therefore, we hypothesize that the learning process is related to the subtle changes in the whole ensemble of different dendritic spine structures, but not at the level of single shape classes.

The source code of our software is freely available for non-commercial use^1^.

## 1 Introduction

Brain plasticity depends on the functional and structural reorganization of synapses. The majority of excitatory synapses are located on dendritic spines, which are small membranous protrusions located on the surface of neuronal dendrites. The important feature of dendritic spines is their structural variability, which ranges from long, filopodia spines to short stubby and mushroom-shaped spines. Dendritic spines are typically built of a head that is connected to the dendrite by a neck. The size of the spine head is proportional to the postsynaptic density area and correlates with postsynaptic receptor content and synaptic strength [12], [21], [30]. The length of the dendritic spine neck is correlated with postsynaptic potential [1], [31]. Thus, dendritic spine shape has been accepted for determining the strength of synaptic connections and is thought to underlie the processes of information coding and memory storage in the brain. Furthermore, alterations in dendritic spine shape, size, and density are associated with a number of brain disorders [4], [8], [9], [13], [14], [22], [25], [28].

The morphology of spines can change in an activity-dependent manner. The structural plasticity of dendritic spines is related to synaptic function, as the morphological modifications of pre-existing spines as well as the formation or loss of synapses accompany learning and memory processes ([32], [33]; for reviews see [3], [7]). The cellular models of synaptic plasticity, *long-term potentiation* (LTP) and *long-term depression* (LTD), associate synaptic strength with spine enlargement and spine shrinkage, respectively [7], [11], [34].

Understanding dendritic spine shape taxonomy and shape transitions upon synaptic potentiation is of great importance. The common approach in morphological analysis of dendritic spines is to categorize spines into subpopulations based on whether they are stubby, mushroom, thin, and filopodia [27]. However, there is a lack of methods allowing for an automatic distinction between dendritic spine subpopulations. To fill this gap, we provide a methodological approach to provide insight into dendritic spine shape taxonomy and transitions in time. Similar to previous works, we potentiated the synapses with LTP stimulation that produces a long-lasting increase in network activity and mimics several aspects of LTP including synaptic receptor incorporation to the dendritic spine membrane. The morphology of single dendritic spines was assessed using time-lapse imaging of living neurons. In the rest of the paper, we refer to a population of spines stimulated by LTP as *ACTIVE*, and the non-treated spines are denoted as *CONTROL*.

In Section 2, we describe the process of data gathering and data representation and the statistical approach to analysis of spine shapes. First, we analyze the basic characteristics of features in populations *ACTIVE* and *CONTROL* and conclude that before a meaningful comparison can be performed, populations need to be normalized. Then, using a split of each population into three subpopulations, growing, not changing and shrinking spines, we compare the relative changes of features across time and note the differences between *ACTIVE* and *CONTROL*. Furthermore, we develop simple but meaningful numerical representations of spines. In Section 3, we provide an approach to dendritic spine taxonomy construction and models of shape transitions together with statistical tests for model comparisons. For taxonomy development, we propose a clustering-based approach that does not depend on subjective decisions of experts and can accommodate arbitrary numerical features. Later, we introduce the corresponding probabilistic model of spine transitions between clusters in time. We also propose a bootstrap-based approach and two statistical tests that are applied for the purpose of the comparison of models built for different populations of spines. Finally, in Section 4, we present our results. We conclude our work in Section 5.

## 2 Data preparation and analysis

In this section, we describe the statistical analyses of the dendritic cell populations *ACTIVE* and *CONTROL*. A comparison of the descriptor distributions showed that initial data preprocessing is necessary, which we performed by carefully choosing subsets of spines from both populations^2^. Further, we introduce the automatic method for dividing each subset into three subpopulations: growing, not changing and shrinking spines. We show how the corresponding subpopulations significantly differ across *ACTIVE* and *CONTROL*. Finally, we introduce the algorithm for spine representation dimensionality reduction.

### 2.1 Data acquisition

Dissociated hippocampal cultures were prepared as described previously in [20]. On the 10th day, *in vitro* cells were transfected using Effectene (Qiagen) according to the manufacturer’s protocol with a plasmid carrying red fluorescence protein under *β*-actin promoter. All the experiments were performed at 19-21 days *in vitro*. Image acquisition was performed using the Leica TCS SP 5 confocal microscope with PL Apo 40 × /1.25 NA oil immersion objective using a 561 *nm* line of diode pumped solid state laser at 10% transmission at a pixel size of 1024 × 1024. Captured cell images consisted of series of z-stacks taken at every 0.4*µm* step. On average, around 14-17 slices (depending on specimen thickness) were taken per stack. The final sampling density was 0.07*µm* per pixel.

The resolution of the confocal microscope along the optical axis (z axis) is three time worse than the resolution along the lateral direction. The majority of observed dendritic spines arise in the lateral direction. Thus, due to limitations of confocal microscopy, it is almost impossible to determine the three-dimensional dendritic spine features. The spines that could be easily distinguished and that protruded in the transverse direction were chosen for analysis. Because of the synaptic scaling, dendritic spine structure and density are modulated with respect to the position along the dendritic tree [17]. To avoid this issue and following the approach by [18], we chose spines that belonged to the secondary dendrites.

The next step of data preparation was to obtain numerical features of the spines. Although many spine extraction methods exist [15], [6], [24], the methods do not prove to be more advantageous than the others. Therefore, we analyzed the images semi-automatically using custom written software [23]. The recorded dendritic spine features were (denoted as *DESCRIPTORS*) length, head width (denote hw), max width location (denote mwl), max width (denote mw), neck width (denote nw), foot, circumference, area, width to length ratio (denote wlr), length to width ratio (denote lwr), and length to area ratio (denote lar). Although researches have not found a consensus yet on which features should be considered, this set covers parameters that are the most often used [18], [29], [31]. The spine length was determined by measuring the curvilinear length along the spine virtual skeleton, which was obtained by fitting the curve (fourth-degree polynomial). The fitting procedure involved searching for a curve along which the integrated fluorescence was at a maximum level. Many spines were distinctly bent such that the distance along a straight line between the tip and the base of the spine underestimates the length of the spine. To define the head width, we used the diameter of the largest spine section that was perpendicular to the virtual skeleton, while the bottom part of the spine (third of the spine length adjacent to the dendrite) was excluded. To define the neck width, we used the thinnest part of the spine between the position of the head-width measurement and the point at which the spine is anchored into the dendrite. Details can be found in [23].

We ended up with two groups of spines, the treatment *ACTIVE* consisting of 433 samples and the control *CONTROL* consisting of 490 samples. For each spine, all of the above 11 features were measured at two different timestamps: *t*_0_ (the time before stimulation) and *t*_1_ (10 minutes after *t*_0_). Researchers showed that after 10 minutes [29], modifications in the spine structure could be already observed, and demonstrated that stimulation causes the cleavage of important adhesion molecules at the dendritic spines [26]. Consequently, by *ACTIVE* (*CONTROL*), we denote all features at all timestamps, and by *ACTIVE*^*x*^, we denote all spines from the *ACTIVE* data, set described only by features at time *t*_*x*_ (similarly, *CONTROL*^*x*^).

### 2.2 Balanced subset selection

In Table 1, we report the mean values for descriptors from *ACTIVE*^0^ and *CONTROL*^0^ populations. We report p-values from two-tailed t-tests for the difference of means between both sets.^3^ We report significant differences for almost all descriptors (only for three features is the p-value above the threshold value *p* > 0.001).

**Table 1.** Differences between *ACTIVE300* and *CONTROL300* at time *t*_0_. Means and p-values from the two-tailed t-test. Descriptor values are measured in *μm* except for the width length ratio and the length width ratio. Significant differences are observed between all descriptor values except for two features.

| feature | *ACTIVE*<sup>0</sup> | *CONTROL*<sup>0</sup> | p-value |
| --- | --- | --- | --- |
| length | 1.268 | 1.539 | 0.000 |
| head width | 0.685 | 0.808 | 0.000 |
| max width location | 0.554 | 0.608 | 0.003 |
| max width | 0.792 | 0.958 | 0.000 |
| width length ratio | 0.667 | 0.657 | 0.721 |
| length width ratio | 2.161 | 2.223 | 0.577 |
| neck width | 0.418 | 0.552 | 0.000 |
| foot | 0.772 | 0.995 | 0.000 |
| circumference | 4.612 | 5.502 | 0.000 |
| area | 0.675 | 0.978 | 0.000 |
| length area ratio | 2.158 | 1.726 | 0.000 |

Such large differences between both sets may influence the statistical analysis of their behavior. Therefore, we decided to preprocess the datasets by excluding some spines, such that the means in the new sets are similar with respect to the statistical test used. Namely, we drew a number of pairs of closest spines, each pair consisting of a spine from the *ACTIVE* set and a spine from the *CONTROL*. The measure of how close the spines are is based on the normalized Euclidean distance^4^ between the vectors of features at time *t*_0_. The pseudo-code for the algorithm is presented in Algorithm S1 in the Supplementary Materials.

In Table 2, we report new statistics on the differences between samples after the 300^5^ closest pairs have been drawn. The same statistical test that was performed before is used here as well. The p-values are significantly higher for all features, and no one feature is significantly different in the two compared groups. We are going to further investigate these new ‘normalized’ sets, denoted as *ACTIVE300* (the 300 closest spines drawn from *ACTIVE*) and *CONTROL300* (the 300 closest spines drawn from *CONTROL*).

**Table 2.** Differences between *ACTIVE300*^0^ and *CONTROL300*^0^ at time *t*_0_. Means and p-values from two-tailed t-tests are shown. Descriptor values are measured in *μm*, except for the width length ratio and length width ratio. No significant differences between any descriptor values are observed for all geometrical features.

| feature | *ACTIVE300*<sup>0</sup> | *CONTROL300*<sup>0</sup> | p-value |
| --- | --- | --- | --- |
| length | 1.240 | 1.276 | 0.416 |
| head width | 0.736 | 0.743 | 0.762 |
| max width location | 0.593 | 0.586 | 0.789 |
| max width | 0.844 | 0.845 | 0.977 |
| width length ratio | 0.702 | 0.688 | 0.630 |
| length width ratio | 1.898 | 1.918 | 0.832 |
| neck width | 0.479 | 0.505 | 0.236 |
| foot | 0.840 | 0.855 | 0.606 |
| circumference | 4.566 | 4.591 | 0.834 |
| area | 0.720 | 0.748 | 0.330 |
| length area ratio | 1.871 | 1.837 | 0.556 |

### 2.3 Division of spines by changing characteristics

In this subsection, we consider relative changes of feature values at times *t*_0_ and *t*_1_. For a fixed feature, relative difference is calculated as: *feature*^rel^ = 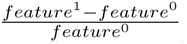.

We consider the relative change in feature length across groups: *ACTIVE300* and *CONTROL300*. Figure 1 shows the relative changes in the feature values for both sets. Note that the *ACTIVE300* population varies more as the corresponding histogram has heavier tails, than *CONTROL300*. We observe that this is the case for all features. We presume that spines from the *ACTIVE300* group compared with *CONTROL300* may exhibit more extreme changes in descriptor values. Therefore, the regions where *ACTIVE* is more frequent than *CONTROL* could possibly be treated as varying. This motivates the following criterion for splitting the spines from both populations into three subgroups: shrinking, not changing and growing. We choose the two separating points defining the three sub-groups such that the differences between the counts of corresponding subgroups from the *ACTIVE300* and *CONTROL300* populations are maximized^6^. The exact method has been shown in Algorithm S2. This criterion assumes that the two groups are of the same size. If they were not, we could easily normalize them by multiplying the appropriate samples from both populations. The summary of the results of the proposed procedure as conducted for feature length is presented in Figure 2.

**Fig. 1.**
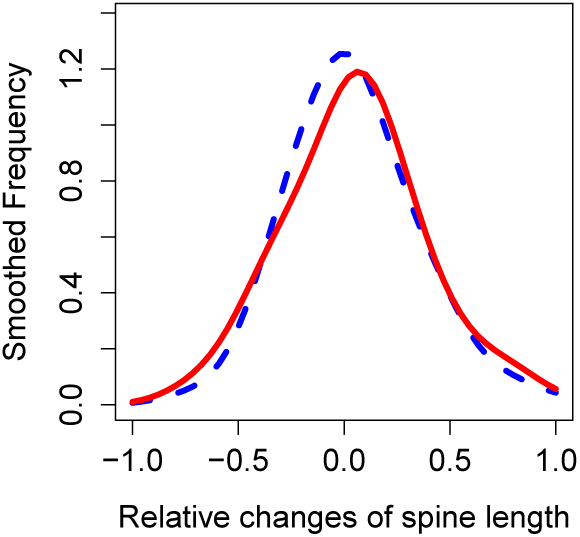
Relative changes in dendritic spine length between time *t*_0_ and *t*_1_ for *ACTIVE300* (solid red) and *CONTROL300* (dashed blue), smoothed using kernel density estimation. We note that *ACTIVE300* varies more, as the corresponding histogram has heavier tails than *CONTROL300*.

**Fig. 2.**
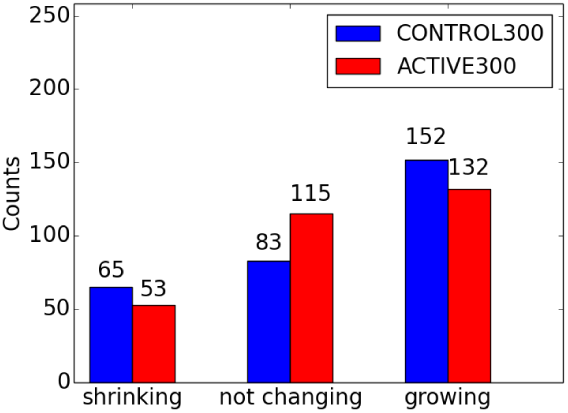
Counts of shrinking, not changing and growing spines from selected populations *ACTIVE300* and *CONTROL300* according to the split based on feature length. No significant differences between the mean values in groups from both populations are observed.

We compared the mean values of features from the subgroups between *ACTIVE300* and *CONTROL300* at time *t*_1_, e.g., to check whether the mean of shortening *ACTIVE300* is different than that of shortening *CONTROL300*. It turns out that there are no significant differences between the populations. In contrast, we applied Pearson’s *χ*^2^ test to check whether the division of the *ACTIVE* spines is similar to the division of *CONTROL* spines. The rationale behind this is that the procedure of dividing spines discriminates between the groups more in terms of the counts obtained than in terms of the means of the subgroups.

The results are reported in Table 3. After the dividing process, Pearson’s *χ*^2^ test was used to evaluate the differences between counts of populations. We notice that all of the obtained p-values, other than those corresponding to the feature foot, are smaller than 5%, which implies that the samples are statistically signstatistically significantly different under the significance level of 5%.

**Table 3.**
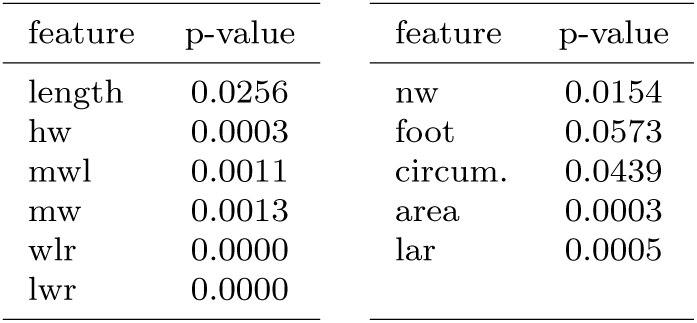
P-values from Pearson’s *X*^2^ test comparing the counts of the growing, not changing and shrinking subgroups between the selected populations *ACTIVE300* and *CONTROL300*. There are significant differences for most descriptors.

### 2.4 Simplification of shape representations

The initial 11 features describing spines can be reduced with the dimensionality reduction technique to render the data representation to be more compact and simple and to filter out the noise. The most popular approach for this purpose is *Principal Component Analysis* (PCA; for details see [10]). We applied PCA to spines from both populations *CONTROL* and *ACTIVE* and for both *t*_0_and *t*_1_. For the first two features (components) in the reduced representation, we cover about 91% of the variance in the data (see Figure S1). The removal of farther features does not reduce the available information by much (only 9% of the variance is lost). The new features are linear combinations of the initial features: *Comp*.1′ = −0.27 · *length -* 0.49 · *lwr* - 0.81 · *circumference* −0.15 · *area*; *Comp*.2′ = −0.17 · *hw* −0.17 · *mw* −0.11 · *wlr* + 0.71 · *lwr* −0.12 · *nw* - 0.12 · *foot* −0.41 · *circumference* −0.21 · *area* + 0.44 · *lar*. We see that *Comp*.1′ is composed mostly of features related to size such as length, circumference, and area. Therefore, this feature can be treated as a generalized size descriptor. Similarly, we can interpret *Comp*.2′ as a generalized contour (shape complexity) descriptor.

The interpretation of the above components as size and contour descriptors allows to construct more meaningful features. The initial features can be directly divided into two sets: *DESCRIPTORS*^*SIZE*^ = {length, circumference, area} (size related features) and *DESCRIPTORS*^*CONTOUR*^ = {hw, foot, mwl, mw, wlr, lwr, lar, nw} (contour related features). Then, PCA is applied separately to each of the sets. Using the first feature from PCA on *DESCRIPTORS*^*SIZE*^ and the first feature from PCA on *DESCRIPTORS*^*CONTOUR*^, 87% of the variance is explained. The loss of the variance compared with PCA computed on all features merged together is equal to 4%. However, the new representation (denoted as *DESCRIPTORS*^*PCA*^) is easy to interpret. New features provide a clear meaning of size and contour complexity and simple form:

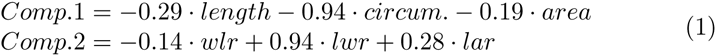

Comparing the loadings (weights) against previous formulas for *Comp*.1′ and *Comp*.2′, we notice that the differences are small, i.e., below 15% in most cases. The most important feature of the size descriptor is the circumference (the highest loading), and the most important feature of the contour descriptor is lwr. Most of the initial features, i.e., hw, foot, mwl, mw, and nw, are not included (they are redundant).

Spine distributions in the new feature space *Comp*.1 × *Comp*.2 are shown in Figure 3. The whole space of features was partitioned into tiles of size 4 × 4, and for each tile, one representative spine (the closest to the tile center) was chosen. We can see how the spine size changes along *Comp*.1 from the smallest on the right side to the biggest one on the left side. Similarly, spine contours change along *Comp*.2, from the simplest on the top to the most complicated on the bottom.

**Fig. 3.**
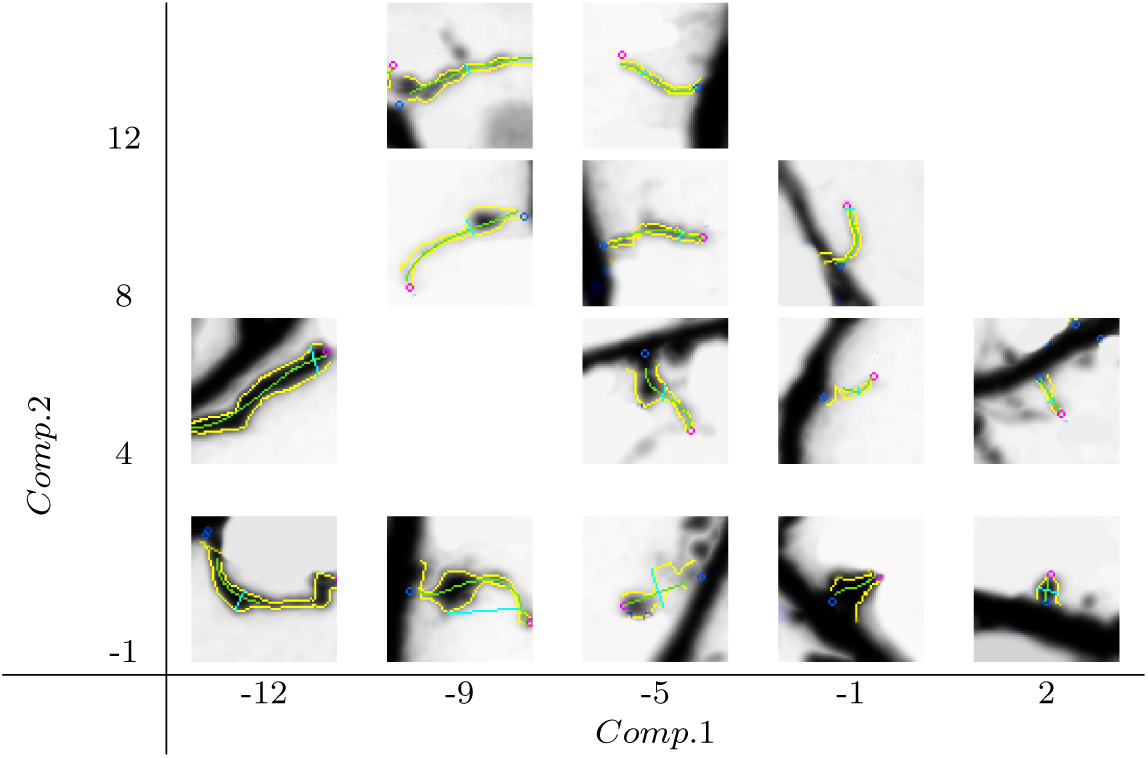
Distribution of spine shapes in space composed of the features *Comp*.1 and *Comp*.2. *Comp*.1 is a generalized size descriptor, and *Comp*.2 is a generalized contour complexity descriptor. Spine sizes change along *Comp*.1 from the smallest on the right side to the biggest on the left side. The spine contour complexity changes along *Comp*.2 from the simplest on the top to the most complicated on the bottom.

## 3 Methods

In this section, we apply two clustering methods to construct the spine shape taxonomy in an unsupervised way. Further, we build the probabilistic model of shape changes in time. Finally, the bootstrap analysis is presented to statistically evaluate differences between both resting and potentiated populations.

### 3.1 Clusters of shapes

Initially, spines are represented in some arbitrary multidimensional space of features, e.g., *DESCRIPTORS*^*PCA*^. Our goal is to obtain a high-level representation that would be both meaningful and simple. Therefore, we propose to apply clustering. Clustering allows for groupings of similar objects (for example, spines) called clusters. Clusters represent possible shapes of spines. The underlying idea is that spines in a cluster have more similar shapes (they are more similar in terms of derived features) among themselves than to spines outside the given cluster. We consider two well-established algorithms, *cmeans* [2] and average-linkage *hierarchical* [19], that represent two main types of clustering, *crisp* and *fuzzy*.

In clustering, each spine *s* is assigned a vector *w*(*s*) = (*w*_1_(*s*),…,*w*_k_(*s*)) of *k* membership weights that are non-negative and sum up to 1. For example, *w*_n_(*s*) is a membership of the spine *s* against the *n*-th cluster. In *crisp* clustering, spines are assigned to exactly one cluster (*w*_*n*_(*s*) = 1 ⇔ *s* assigned to n-th cluster; 0 otherwise). In *fuzzy* clustering, weights can be arbitrary real numbers between 0 and 1. Additionally, weights can be interpreted as probabilities, e.g., *w*_*n*_(*s*) can be interpreted as the probability that spine *s* belongs to the *n*-th cluster.

To obtain a taxonomy of shapes that would describe spines in both time points equally well, we applied clustering to data *ACTIVE*∪ *CONTROL* from both time points *t*_0_ and *t*_1_. Consequently, each spine was included twice and assigned two vectors of weights. Spine *s* at time *t*_0_ is assigned the vector *w*^0^(*s*) and at time *t*_1_ the vector *w*^1^(*s*). We denote 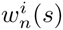 = *P*^*i*^(*s* ∈ *C*_*n*_) as the probability that spine *s* belongs to the cluster *n* at time *t*_*i*_.

The prediction of weights of a new spine *s* (not in the training data) is not always obvious. For *hierarchical* clustering, we used a 1-nn classifier, i.e., we search for the most similar sample vector *s′* from the training data and assign *w*(*s*) = *w*(*s*′). In *cmeans* clustering, the prediction of weights of a new spine *s* is more straightforward. Each spine, whether from the training data or not, has weights assigned according to the same explicit formula.

The above clustering algorithms have either one (*hierarchical*) or two (*cmeans*) parameters: *k* - number of clusters and *m* - fuzzifier (informs about clusters fuzziness). Large *m* results in smaller weights and more fuzzy clusters. For small *m*, e.g., *m* = 1, we obtain results close to *crisp* clustering. Consequently, low values of both *k* and *m* are preferred. Although these parameters can be selected in many ways, we decided to use *Within Cluster Sum of Squares* (*WSS*), as it has several good properties, i.e., simple meaning, applicability to both crisp and fuzzy cases, and the same behavior no matter what data and what clustering algorithm are used (it decreases when *k* increases and when *m* decreases). For balance between the number of clusters, data fitness values of *k* and *m* at ‘knee point’ (the point where *WSS* plot bends the most) should be selected. The definition of *WSS* is as follows: *WSS* = Σ_n=1‥k_ Σ_s_ *w*_*n*_(*s*)(**s** - **c**_**n**_)^2^ where **c**_**n**_ = 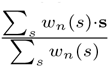, where s stands for the vector of features assigned to object *s* and **c**_**n**_ is *n*-th cluster centroid.

### 3.2 Shape transition model

#### Assumptions and brief description

Researchers showed that the initial dendritic spine morphology may influence how this structure will change upon specific treatment [29], [16], e.g., induction of long-term potentiation. Therefore, we assume that changes of spines depend on their initial shapes and that each spine follows patterns of behavior highly correlated with its initial shape. We introduce the novel probabilistic model of behavior that relies on these principles.

We represent spines as combinations of shapes and spine changes with combinations of behavior patterns. Shape combinations are represented by weights of shape clusters *w*_*n*_(*s*). Combinations of behavior patterns are represented with probabilities *P*(*C*_n_ → *C*_*m*_|*C*_*n*_), or the probability that the shape represented by cluster *C*_*n*_ will change into the shape represented by cluster *C*_*m*_ when *t*_0_ → *t*_1_. Probabilities *P* can be stored in a *k* × *k* matrix called *transition matrix*, where rows are enumerated with *n* and columns with *m*. An even more convenient representation of the same information is a graph, where nodes represent shape clusters and edges are labeled with probabilities, denoted as a *transition graph*.

#### Probability estimation

In the *crisp*, e.g., *hierarchical* model of shapes, we can estimate the probability *P* as follows: *P*_*crisp*_(*C*_*n*_ → *C*_*m*_|*C*_*n*_) = 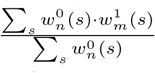. In the denominator, we have a number of spines that belong to cluster *C*_*n*_ in time *t*_0_ (normalizer). In the denominator, there is a number of spines that belong to cluster *C*_*n*_ in time *t*_0_ and to cluster *C*_*m*_ in time *t*_1_ (recall that only for one *n* in 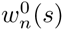 and for one *m* in a 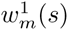, the values are ones; elsewhere, they are zeros). With such a computation, we consider how many spines moved from shape cluster *C*_*n*_ to *C*_*m*_ and normalize it by the number of all spines in the initial cluster *C*_*n*_.

There are arbitrarily many generalizations that are consistent with the above crisp derivation for the fuzzy model, e.g., *cmeans* model, i.e., 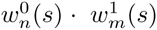 can be reformulated in many ways without changing the values of *P*_*crisp*_, e.g., as min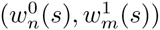. We suggest using the generalization for which the model minimizes the error of behavior prediction of a spine *s* when *t*_0_ → *t*_1_. The probability that spine *s* in time *t*_1_ will be in cluster *C*_*m*_ for our linear model is given according to the law of total probability as follows:

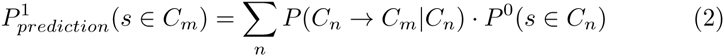

The overall prediction error can be computed as a sum of squared differences between predicted 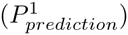 and derived probabilities (*P*^1^):

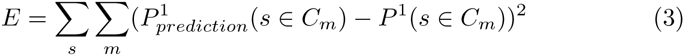

where for each spine *s* in the data, we compare the membership for cluster *C*_*m*_ at time *t*_1_ with the prediction of the model. The problem can be now formulated as an optimization task where we search for probabilities *P*(*C*_*n*_ → *C*_*m*_|*C*_*n*_) that minimize the overall prediction error *E*:

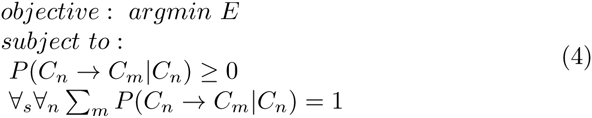

The above derivations can be easily represented in matrix form, and the above optimization problem is an example of a standard quadratic programming optimization task with constraints. Details are presented in Section S7 in Supplemental Materials.

#### Parameter reliability

To derive information on the reliability of the obtained probabilities, we use the following bootstrap-based procedure. We generate *R* = 1000 new populations sampled with replacement from the original population. For each new population, we calculate all the probabilities again. The average squared differences between probabilities for new populations and the original populations are used as the estimates of parameter errors. Formally, the error of the probability *P*(*C*_*n*_ → *C*_*m*_|*C*_*n*_) is calculated as:

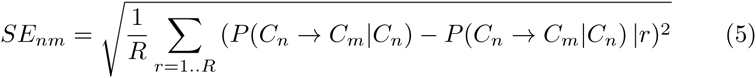

where *P*(*C*_*n*_ → *C*_*m*_|*C*_*n*_)|*r* denotes the probability calculated for the *r*-th bootstrap population.

### 3.3 Comparison of models

Bootstrap Hypothesis Testing [5] is a method of testing statistical hypotheses. To apply the method, one has to first modify the testing sample so that the null hypothesis is satisfied. Subsequently, a large number of bootstrap samples is drawn from such a modified sample. Finally, for the fixed statistic of interest, one must evaluate how extreme the value of the statistic is for the original sample compared with the values obtained for the drawn bootstrap samples.

This general rule in our case proceeds as follows. We take the two groups *ACTIVE300* and *CONTROL300* and join them into one group *ACTIVE300*∪ *CONTROL300*. At each iteration of bootstrap sampling, two new groups are drawn from the joint dataset. This way, the null hypothesis of a common distribution for both groups is satisfied. Next, for each bootstrap, the sample clusters and *Shape Transition Model* are constructed for both groups. Then, the test statistic is computed. Finally, the statistic is computed on models built for original groups and compared with the bootstrap sampling results.

#### Comparison of changes in cluster distributions

We cluster spines according to their shapes (see Section 3.1). As a result, for each spine *s* at *t*_0_ and *t*_1_, we obtain the set of weights representing a mixture of shapes. Then, we derive the overall distribution (total weights) of shapes (by shapes, we mean shape clusters) at both *t*_0_ and *t*_1_. The *n*-th cluster total weight (in case of *crisp*, e.g., *hierarchical* clustering, it is equivalent to number of spines) in *t*_0_ is equal to 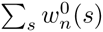 and in *t*_1_ is equal to 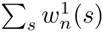. Consequently, the relative change in the *n*-th cluster weight between *t*_0_ and *t*_1_ for population *G* can be computed as follows: *C*_*n*_(*G*) = 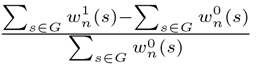. The statistic that measures the difference between relative changes in distributions of shapes for populations *G*_1_, *G*_2_ can be now defined as:

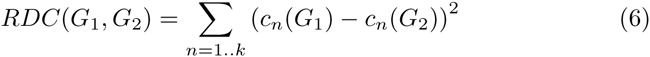

#### Comparison of transition matrices

By applying the *Shape Transition Model* (see Section 3.2), we construct two Markov matrices (*transition matrices*) describing transitions for both populations. To check how similar the matrices are, we decided to apply bootstrap hypothesis testing. For comparing the matrices, we use the sum of squared differences between corresponding cells from the two matrices:

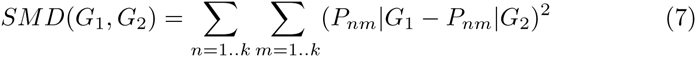

where *G*_1_,*G*_2_ are populations, e.g., *ACTIVE300*, *CONTROL300*, to be compared. *P*_*nm*_|*G*_i_ ≡ *P*(*C*_*n*_ → *C*_*m*_|*C*_*n*_)|*G*_*i*_ stands for the value of a cell in the *n*-th row and in the *m*-th column of the transition matrix *P* built with data from population *G*_*i*_.

## 4 Results

To obtain the taxonomy of spine shapes, we applied *cmeans* and *hierarchical* clustering to *ACTIVE*∪ *CONTROL* for *t*_0_ and *t*_1_. To select the proper values of the parameters, we used *WSS* plots with ‘knee’ shapes (see Figure S3). We obtained *k* = 10 for *hierarchical* and *k* = 8, *m* = 4 for *cmeans* clustering. According to *WSS* measures, these values ensure a good balance between the complexity of results, i.e., the number of clusters and quality of cluster fitness.

Figure 4(a) presents the results of *hierarchical* clustering calculated for *ACTIVE*∪ *CONTROL* according to the procedure described in Section 2.4. Each spine is represented with a single point, and the colors represent the cluster memberships. For each cluster, we identified three representative spines lying nearest to the cluster center. Representative spines are shown in Figure 4(b). Obtained clusters express the universal taxonomy of shapes that will be later employed for the computation of *Shape Transition Model* for *ACTIVE*, *CONTROL*, *ACTIVE300* and *CONTROL300*. Representative spines can be used for visual inspection and biological interpretation.

**Fig. 4.**
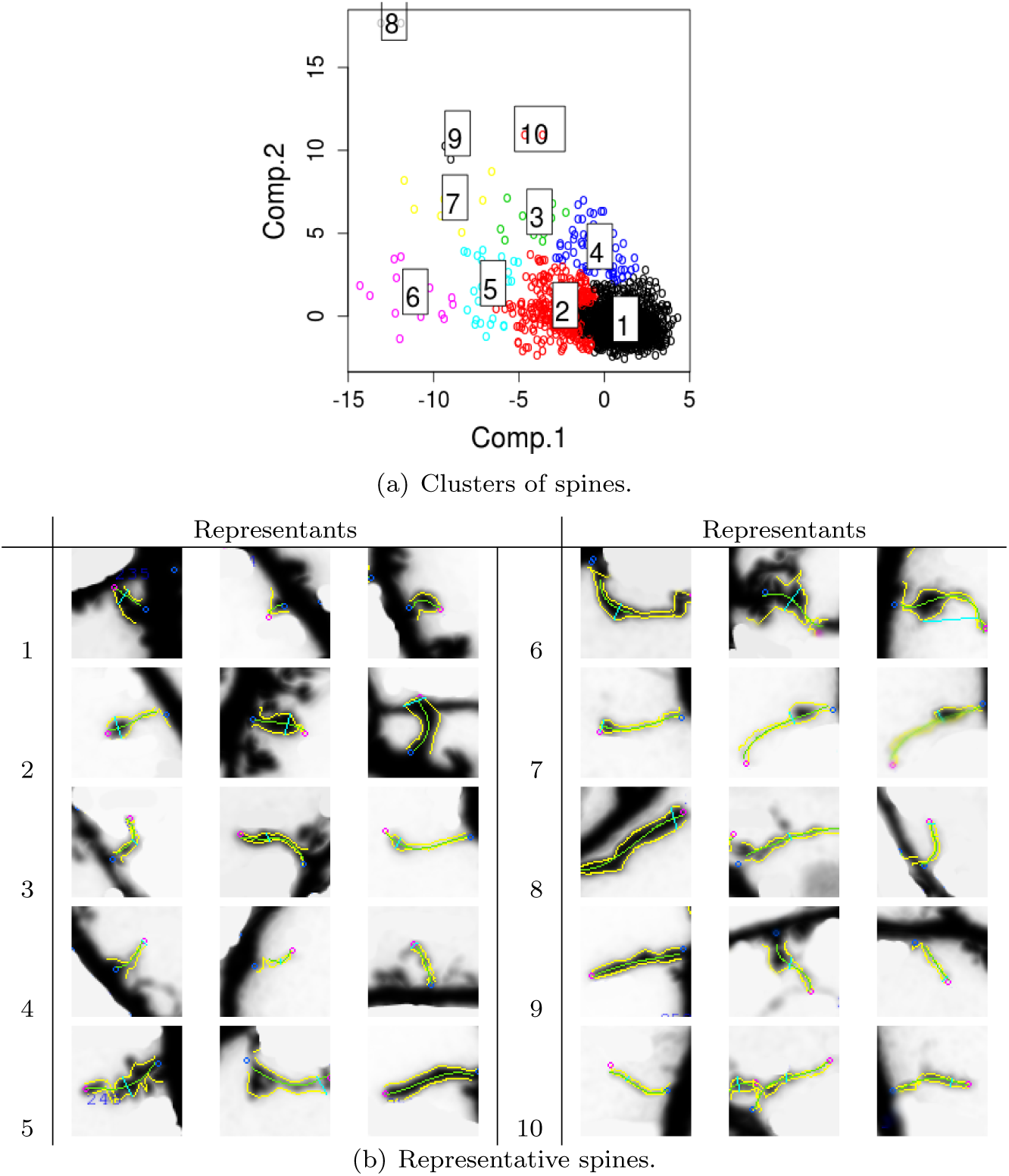
*Hierarchical* clustering and representative spines obtained for *ACTIVE* ∪ *CONTROL*. The presented clusters represent the universal taxonomy of spine shapes. For each cluster, we present three spines that are nearest to the cluster center. Representative spines facilitate visual aid for interpretation purposes.

Apart from *hierarchical* clustering, we also consider *cmeans* clustering. Table 4 presents the comparison of the prediction error *E* for both methods. Values were obtained using 10-fold cross-validation. Numbers from the same column but in different rows should not be compared. Different clustering methods result in different shape clusters that have different members and thus are incomparable. Although errors *E* for different methods have different ranges and cannot be compared, different models with the same method can be compared.

**Table 4.** Prediction error *E* for various models and clustering methods. Values were obtained using 10-fold cross-validation on *ACTIVE*∪ *CONTROL*. Values in columns should not be compared. For both clustering methods, *Shape Transition Model* performs better than the baseline.

| Clustering method | *Shape Transition Model* | Majority vote | No transitions | Random transitions |
| --- | --- | --- | --- | --- |
| *hierarchical* | 0.266 +/-<br>0.147 | 0.395 +/-<br>0.237 | 0.433 +/-<br>0.242 | 0.997 +/-<br>0.124 |
| *cmeans* | 0.024 +/-<br>0.004 | 0.853 +/-<br>0.029 | 0.037 +/-<br>0.007 | 0.054 +/-<br>0.012 |

The *Shape Transition* Model is compared with three baselines. The first baseline is the majority vote model, where all spines from a particular cluster move to a single destination cluster that is selected as the most popular choice. The second baseline is the model, where we assume that all spines remain in the initial clusters, i.e., weights in *t*_1_ are the same as in *t*_0_. Finally, the third baseline assumes random values for probability *P*. For both clustering methods, the *Shape Transition* Model has the smallest error *E* and predicts spine behavior the best.

*Transition graphs* of the *Shape Transition Model* for *ACTIVE* and *CONTROL* are shown in Figures 5(a) and 5(b). Each cluster of shapes is represented by an oval. Initial sizes, i.e., weights of clusters (for *hierarchical* clustering equivalent to number of spines), are listed. Edges representing transitions are labeled with probability P. They are filtered out, and only transitions (probabilities) of greater than 20% of the initial weight are visible.

**Fig. 5.**
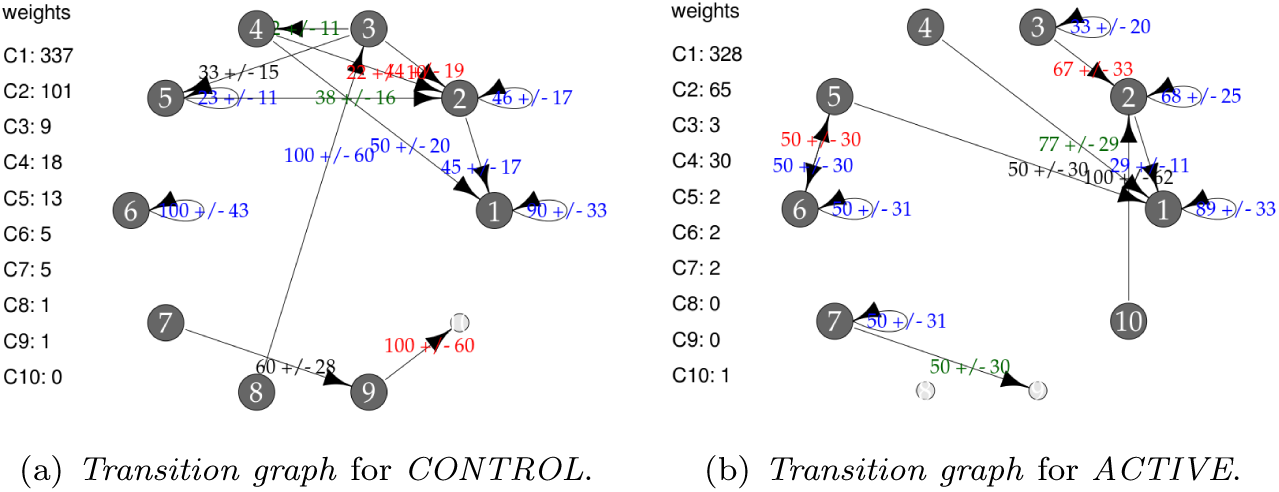
*Transition graphs* for *hierarchical* clustering. For each cluster, the initial weight (number of spines in the cluster) is presented. Only transitions (probabilities) of values higher than 20% are shown. Only clusters 1-5 are well represented in the data. Transitions for the remaining clusters are uncertain.

Only five clusters (numbers 1-5) are well represented in the data. Clusters 1, 2 and 4 are the most dense. Clusters 3 and 5 are interpreted as peripheral. Finally, clusters 6-10 have only a few spines. For transitions from clusters 6-10, high errors were obtained. For example, *SE* for *P*(*C*_9_ → *C*_10_|*C*_9_) is equal to 66%. Conclusions concerning clusters 6-10 are not reliable. Analogous plots of clustering results and *transition graphs* for *cmeans* are presented in Figures S4, 5(a) and 5(b).

Graphs presented in Figures 5(a) and 5(b) should not be compared because they are computed for populations of different characteristic at *t*_0_. Alternatively, Figure 6 presents the comparison of *transition graphs* for *CONTROL300* and *ACTIVE300* for *hierarchical* clustering (exact values of the probabilities can be found in Table S1). A similar analysis for *cmeans* is presented in Figure S6, and the values of the transitions in percents can be found in Table S2.

**Fig. 6.**
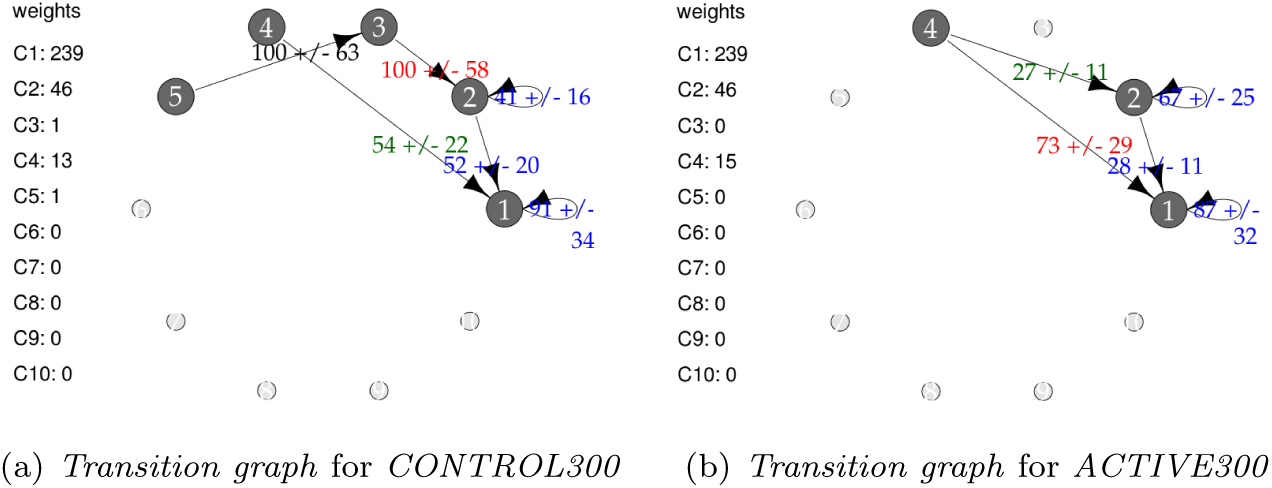
Comparison of the *transition graphs* for balanced subpopulations and *hierarchical* clustering. For each cluster, the initial weight (number of spines in the cluster) is presented. Values are given in percents. Only transitions (probabilities) of values higher than 20% are shown. Differences in transitions between graphs are observed, but because of high uncertainties, none of them is significant.

In the case of the *CONTROL300* and *ACTIVE300* subsets (Figure 6), only clusters 1, 2 and 4 contain enough spines to produce credible conclusions. For *CONTROL300*, cluster 1 has slightly stronger inertia than for *ACTIVE300* (91% vs. 87% spines remained in the same cluster). For cluster 2, the situation is the opposite: 41% of spines from cluster 2 for *CONTROL300* remain in cluster 2 compared with 67% for *ACTIVE300*. For both populations, a large transition of spines from cluster 2 to cluster 1 is observable. However, for *CONTROL300*, it is present for 52% of the spines, whereas for *ACTIVE300*, it is present only for 28%. Another difference is visible for transitions from cluster 4. For *CONTROL300*, 73% of spines move to cluster 1 and 27% to cluster 2. For *ACTIVE300*, only 54% of spines move to cluster 1, and the rest move to clusters 2-5. Unfortunately, none of the observed differences is significant when the errors are taken into consideration. Therefore, to identify such differences, the models must be compared as a whole.

Table 5 presents p-values of *RDC* and *SM D* statistics used for a comparison of models for *ACTIVE300* and *CONTROL300*. Results below 0.05 are marked in bold font. Detailed plots of the statistical distributions using kernel estimation are shown in Figures S7 and S8. For *hierarchical* clustering, only *SM D* shows a significant difference between *ACTIVE300* and *CONTROL300*. This statistic compares transitions of spines between shapes, which is well captured by *hierarchical* clustering. The *RDC* statistic relies only on changes of distributions, and *hierarchical* clustering enforces that each spine belongs to only one shape cluster at the particular time point, which may noticeably affect the overall distributions. In contrast, distributions are well captured by *cmeans* clustering, where each spine is an arbitrary mixture of shapes and *RDC* shows a significant difference. Different clustering methods are sensitive to different properties of the data. The selection of the right clustering method and appropriate test depends on the characteristic of the data that is of interest to the researcher.

**Table 5.**
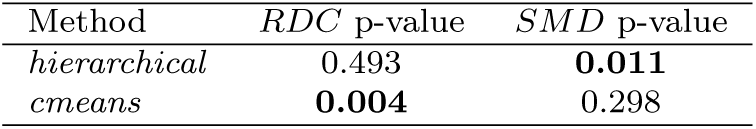
P-values of *RDC* and *SM D* statistics with bootstrap tests used to compare balanced subpopulations *ACTIVE300* and *CONTROL300* for various clustering methods. Differences that are statistically significant are shown in bold font.

| Method | *RDC* p-value | *SMD* p-value |
| --- | --- | --- |
| *hierarchical* | 0.493 | **0.011** |
| *cmeans* | **0.004** | 0.298 |

## 5 Discussion and conclusions

The majority of excitatory synapses in the brain are located on dendritic spines. These highly dynamic and plastic structures undergo constant morphological changes in different physiological and pathological processes [11]. The structure of the dendritic spines is tightly correlated with their function and reflects the synapse properties. Synapse strengthening or weakening along with dendritic spine formation and elimination assure correct processing and storage of the incoming information in the neuronal network. This plastic nature of the dendritic spines allows them to undergo activity-dependent structural modifications, which are thought to underlie learning and memory formation. At the cellular level, the most extensively studied aspect of this phenomena is related to dendritic spine enlargement in response to stimulation.

In this study, we explored the impact of the externally applied stimulation on the dendritic spine structural dynamics. We applied statistical tests and examined a population consisting of 923 dendritic spines. We used two dissociated neuronal cell cultures and compared dendritic spine volume and shape changes between two populations at two different states, unstimulated (*CONTROL*) and LTP-stimulated (*ACTIVE*), and at two time points (with a 10-minute time interval). We preprocessed the datasets and reduced the dendritic spine number to 300 for each analyzed group. We introduced an automatic way of splitting the populations into growing, not changing, and shrinking spines and showed that the two-dimensional descriptors of dendritic spine change differently between the corresponding populations in a significant manner.

The obtained results show that changes in the dendritic spine shape and size are associated with neuronal cell activation upon stimulation. By employing statistical analysis, we confirmed that neuronal activity influences the overall composition of the dendritic spine population. Additionally, we provided a probabilistic model for dendritic spine population dynamics. First, the resting state model was constructed (Figure 5(a)). Then, the probabilistic null model for active neurons was built (Figure 5(b)). We showed that LTP treatment induced transition of filopodia-like spines (cluster **4**) into mushroom-shaped spines (cluster **2**). For the first time, we provided the exact transition probabilities for this morphological transformation (from cluster **4** to cluster **2**, the transition probability was found to be 0.27 ± 0.11). Our result supports the previous studies [29] who report chemical LTP-induced spine enlargement in dissociated cultures.

Finally, we compared models for balanced populations (Figure 6). We found differences between active and non-active neurons. Unfortunately, none of the observed differences between the models was significant when particular transitions between shape clusters were considered. Large errors predominated the differences between values in cells of appropriate transition matrices. However, statistically significant differences were detected when whole models of populations were compared. Different clustering algorithms showed statistically significant differences between the two analyzed groups (*ACTIVE300*, *CONTROL300*). Crisp clustering captured the difference in shapes transitions well, whereas fuzzy clustering captured the difference in changes of shape cluster distributions.

We hypothesize that biological information is not stored in the specific spines shapes or sizes; rather, it is related to the dynamical changes at the population level. The subtle changes in the relative number of dendritic spines for each structural type, matching different shapes or volumes, could be responsible for forming or rearranging information-processing pathways in neuronal networks. According to our model, the dynamics of the entire dendritic spine groups, not the individual entities, are at the center of this cellular phenomena. This may serve to better understand the complex mechanism of information processing in the brain.

## Acknowledgements

The authors are grateful to Prof. Jakub Wlodarczyk, Prof. Grzegorz Wilczynski and Prof. Subhadip Basu for extensive discussions on the information processing problem in neuronal systems and especially to Blazej Ruszczycki for preparation of the segmentation software used for acquiring the morphological descriptors. Some calculations were performed at the Interdisciplinary Centre for Mathematical and Computational Modelling, University of Warsaw, grant No. G49-19. The data used in the experimental part of the study were gathered in the Laboratory of Cell Biophysics at Nencki Institute of Experimental Biology under the supervision of Prof. Jakub Wlodarczyk.

## Computational approach to dendritic spine taxonomy and shape transition analysis

Supplemental Materials

Tomasz Kusmierczyk^1^, Michal Lukasik^2^, Marta Magnowska^3^, Matylda Roszkowska ^3^, Grzegorz Bokota^4^, Dariusz Plewczynski^4,5^

^1^ Department of Computer and Information Science, Norwegian University of Science and Technology, ^2^ Department of Computer Science, University of Sheffield, ^3^ Nencki Institute of Experimental Biology, Polish Academy of Sciences, ^4^ Centre of New Technologies, University of Warsaw, Poland, ^5^ Faculty of Pharmacy, Medical University of Warsaw, Poland

## S6 Algorithms for subset selection and group division

Large differences between the sets *ACTIVE* and *CONTROL* may influence the statistical analysis of their behavior. Therefore, we decided to preprocess the datasets by excluding some spines, such that the means in the new sets are close with respect to the statistical test used. Below we show pseudocode for algorithm, where subsets of spines are selected, forming new sets for further analysis.

### Algorithm S1 SUBSET-SELECTION

Input:

Lists of spines: *ACTIVE* and *CONTROL*

A function of state of all variables: *STOP CONDITION*

Output:

Lists of spines: *ACTIVESUBSET* and *CONTROLSUBSET*

1: Normalize each feature of *ACTIVE* and *CONTROL* by subtracting the common mean and dividing by the common standard deviation,

2: Initialize *ACTIVESUBSET* and *CONTROLSUBSET* to empty lists,

3: **while** *STOP CONDITION* is not satisfied do

4: draw the pair of spines *x*1 ∈ *ACTIVE* and *x*2 ∈ *CONTROL* of the smallest euclidean distance

5: move *x*1 and *x*2 from their lists respectively to *ACTIVESUBSET* and *CONTROLSUBSET*

6: **end while**

7: **return** *ACTIVESUBSET* and *CONTROLSUBSET*

We choose the two separating points defining the three sub-groups such that the differences between the counts of corresponding subgroups from the *ACTIVE300* and *CONTROL300* populations are maximized^7^. The exact method has been shown in Algorithm S2. This criterion assumes that the two groups are of the same size. If they were not, we could easily normalize them by multiplying the appropriate samples from both populations. See Figure 2 for summary of the results.

Let 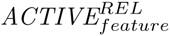 denote the representation of *ACTIVE* with relative differences of feature feature. Similarly for 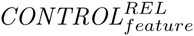 and subsets 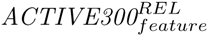 and 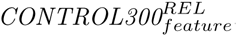.

### Algorithm S2 SEPARATING-POINTS

Input:

Lists of feature values: 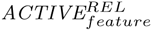 and 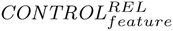, for a fixed feature Output:

A value: *SPLITPOINT*

1: Sort 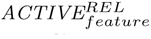 and 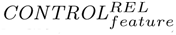 increasingly,

2: Initialize counters: *ACTIVECNT* = 0 and *CONTROLCNT* = 0,

3: Initialize information on the splitting point: *SPLITV AL* = 0, *SPLITPOINT* = *NULL*

4: **while** at least one list: 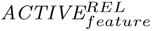 or 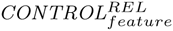 is non-empty **do**

5: *x* = smaller of the 2 elements at the front of the lists: 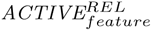 and 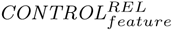,

6: delete all occurrences of *x* from 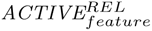 and add the number of its occurrences to *ACTIVECNT*

7: delete all occurrences of *x* from 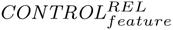 and add the number of its occurrences to *CONTROLCNT*

8: **if** *SPLITV AL* < *ACTIVECNT* - *CONTROLCNT* **then**

9: *SPLITV AL* = *ACTIVECNT* - *CONTROLCNT*

10: *SPLITPOINT* = *x*

11: **end if**

12: **end while**

13: **return**SPLITPOINT

## S7 Matrix formulation of Shape Transition Model

*Shape Transition Model* can be represented in the matrix form where:

– *W* ^*i*^ - *N* × *k* matrix of weights where each row represents a single spine at time *t*_*i*_

– *P* - *k* × *k* matrix of transition probabilities *P*(*C*_*n*_ → *C*_*m*_|*C*_*n*_) indexed by *n* and *m*.

Predictions of the model can be calculated as follows:

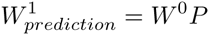

Prediction error can be calculated as follows 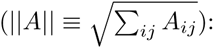

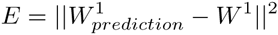

The optimization problem is given by: (1_*k*_ - *k*-element vertical vector of ones):

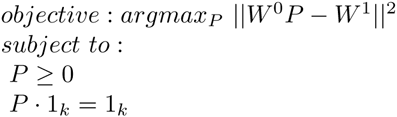

and can be transformed to the standard quadratic programming form:

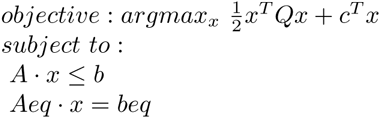

where:
– *x* = *x*(*P*) is a vector of length *k*^2^
– *Q* = *Q*(*W* ^*i*^, *W* ^*j*^), *A* = *A*(*W* ^*i*^, *W* ^*j*^), *Aeq* = *Aeq*(*W* ^*i*^, *W* ^*j*^) are matrices of size *k*^2^ × *k*^2^
– *c* = *c*(*W* ^*i*^, *W* ^*j*^), *b* = *b*(*W* ^*i*^, *W* ^*j*^), *beq* = *beq*(*W* ^*i*^, *W* ^*j*^) are vectors of length *k*^2^

## S8 Supplemental figures and tables

**Fig. S1.**
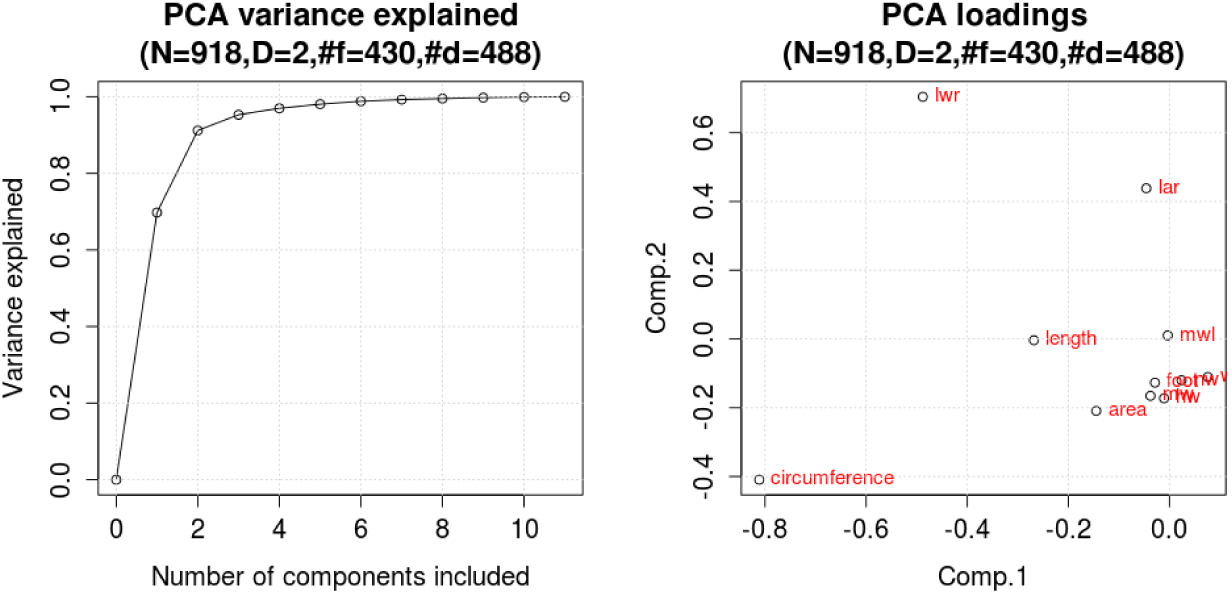
Proportion of the explained variance for different numbers of components (left) and loadings (weights) for two of the most important components (right). PCA was calculated on *DESCRIPTORS* of *CONTROL* ∪ *ACTIVE* data. For two features (components) about 91% of the variance is explained. We see that *Comp*.1′ is composed mostly of features related to size such as length, circumference, and area. Therefore, this feature can be treated as a generalized size descriptor. Similarly, we can interpret *Comp*.2′ as a generalized contour (shape complexity) descriptor.

**Fig. S2.**
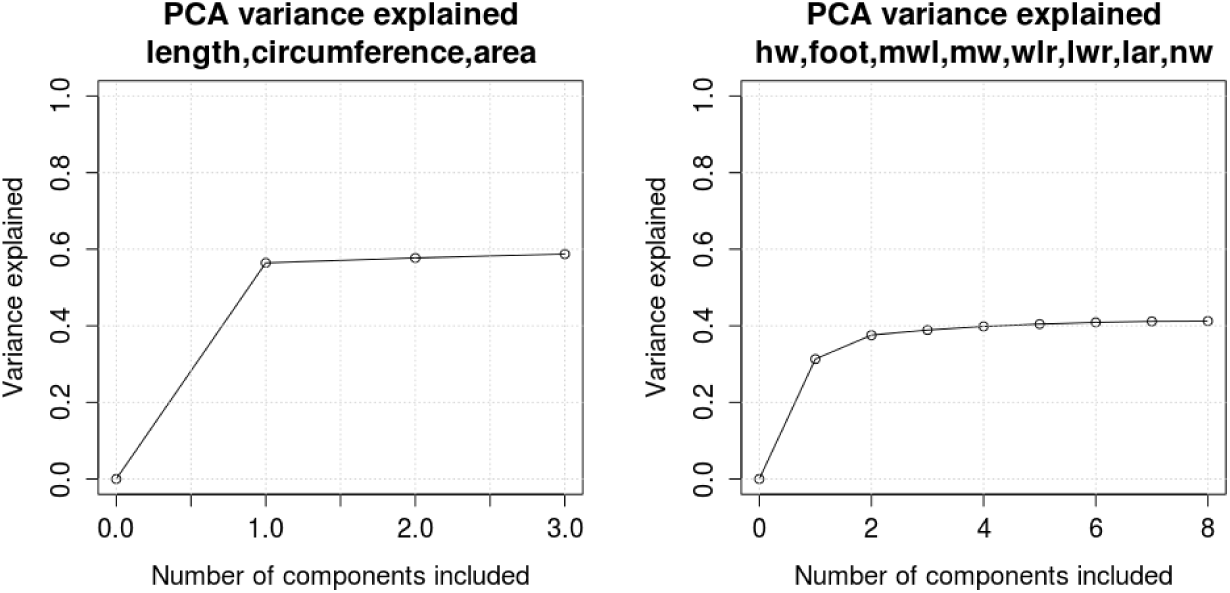
Proportion of the explained variance for PCA on components (features) describing size (left) and contour (right). PCA was calculated separately on *DESCRIPTORS*^*SIZE*^ = {length, circumference, area} (size related features) and on *DESCRIPTORS*^*CONTOUR*^ = {hw, foot, mwl, mw, wlr, lwr, lar, nw} (contour complexity related features) of *CONTROL*∪ *ACTIVE* data. Using the first feature from PCA on *DESCRIPTORS*^*SIZE*^ and the first feature from PCA on *DESCRIPTORS*^*CONTOUR*^ 87% of the variance is explained.

**Table S1.** *Transition matrices t*_0_ → *t*_10_ for *CONTROL300* and *ACTIVE300* for *hierarchical* clustering. Values are denoted in percents, *SE* in brackets, source clusters in rows, and destination clusters in columns. Only clusters 1, 2 and 4 contain enough spines to produce credible conclusions. According to estimated errors, transitions observed for other cases are not meaningful.

| *CONTROL300* |  |  |  |  |  |  |  |
| --- | --- | --- | --- | --- | --- | --- | --- |
| Cluster | 1 | 2 | 3 | 4 | 5 | 6 | 7-10 |
| 1 | 91 (34) | 6 (2) | 0 | 2 (1) | 0 | 0 | 0 |
| 2 | 52 (20) | 41 (16) | 0 | 2 (1) | 2 (1) | 2 (1) | 0 |
| 3 | 0 | 100 (58) | 0 | 0 | 0 | 0 | 0 |
| 4 | 54 (22) | 15 (8) | 8 (5) | 15 (8) | 8 (5) | 0 | 0 |
| 5 | 0 | 0 | 100 (63) | 0 | 0 | 0 | 0 |
| 6 | 0 | 0 | 0 | 0 | 0 | 0 | 0 |
| 7-10 | 0 | 0 | 0 | 0 | 0 | 0 | 0 |

| *ACTIVE300* |  |  |  |  |  |  |  |
| --- | --- | --- | --- | --- | --- | --- | --- |
| Cluster | 1 | 2 | 3 | 4 | 5 | 6 | 7-10 |
| 1 | 87 (32) | 12 (4) | 0 | 0 | 0 | 0 | 0 |
| 2 | 28 (11) | 67 (25) | 0 | 2 (1) | 2 (1) | 0 | 0 |
| 3 | 0 | 0 | 0 | 0 | 0 | 0 | 0 |
| 4 | 73 (29) | 27 (11) | 0 | 0 | 0 | 0 | 0 |
| 5 | 0 | 0 | 0 | 0 | 0 | 0 | 0 |
| 6 | 0 | 0 | 0 | 0 | 0 | 0 | 0 |
| 7-10 | 0 | 0 | 0 | 0 | 0 | 0 | 0 |

**Fig. S3.**
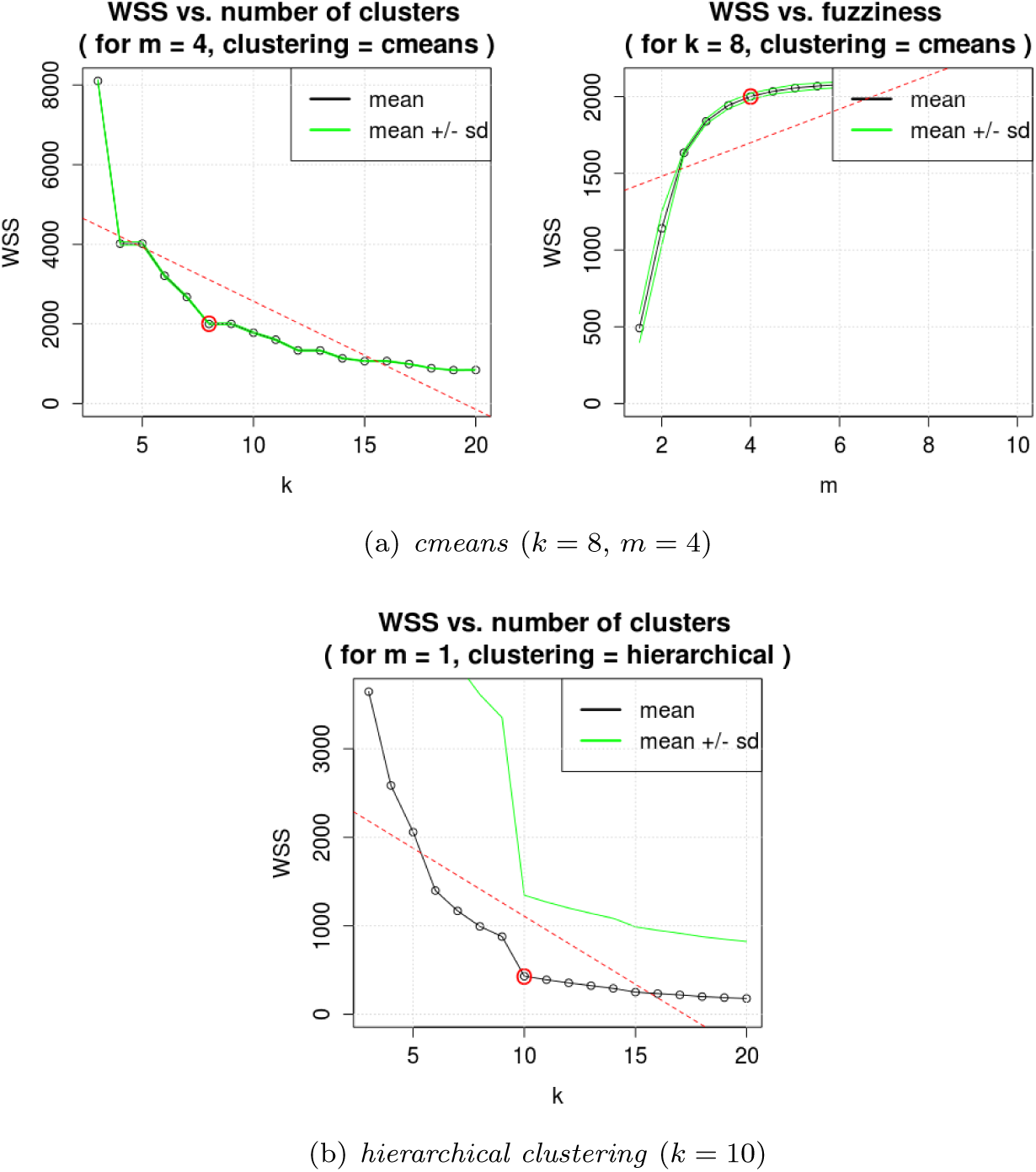
Parameters selection for shape clustering methods (*WSS* plots for *ACTIVE* ∪ *CONTROL*). Red circles mark the selected values in ‘knee points’.

**Table S2.** Transition matrices *t*_0_ → *t*_10_ for *CONTROL300* and *ACTIVE300* for *cmeans* clustering. Values are denoted in percents, *SE* in brackets, source clusters in rows, and destination clusters in columns. In contrast to *hierarchical* clustering case all clusters contain nonneglible amounts of spines.

| *CONTROL300* |  |  |  |  |  |  |  |  |
| --- | --- | --- | --- | --- | --- | --- | --- | --- |
| Cluster | 1 | 2 | 3 | 4 | 5 | 6 | 7 | 8 |
| 1 | 43 (18) | 5 (4) | 0 (1) | 4 (3) | 22 (11) | 0 (4) | 0 (3) | 25 (12) |
| 2 | 2 (1) | 17 (6) | 13 (5) | 10 (4) | 3 (2) | 21 (8) | 28 (10) | 6 (3) |
| 3 | 0 (0) | 5 (2) | 50 (19) | 0 (1) | 0 (0) | 15 (6) | 29 (12) | 1 (1) |
| 4 | 1 (1) | 19 (7) | 5 (2) | 35 (13) | 10 (4) | 14 (7) | 4 (2) | 12 (5) |
| 5 | 11 (5) | 11 (5) | 3 (2) | 23 (9) | 27 (12) | 3 (3) | 2 (2) | 20 (8) |
| 6 | 2 (1) | 20 (8) | 10 (4) | 8 (4) | 5 (3) | 40 (14) | 15 (6) | 0 (1) |
| 7 | 5 (2) | 15 (6) | 19 (8) | 17 (7) | 4 (2) | 10 (4) | 21 (8) | 9 (4) |
| 8 | 4 (3) | 19 (8) | 2 (2) | 14 (6) | 17 (8) | 9 (5) | 11 (6) | 24 (10) |

| *ACTIVE300* |  |  |  |  |  |  |  |  |
| --- | --- | --- | --- | --- | --- | --- | --- | --- |
| Cluster | 1 | 2 | 3 | 4 | 5 | 6 | 7 | 8 |
| 1 | 33 (15) | 19 (9) | 7 (4) | 0 (4) | 11 (7) | 0 (1) | 12 (6) | 19 (9) |
| 2 | 2 (1) | 23 (9) | 5 (3) | 25 (10) | 5 (3) | 15 (8) | 8 (4) | 16 (7) |
| 3 | 1 (1) | 13 (5) | 40 (15) | 2 (1) | 0 (0) | 13 (5) | 27 (10) | 3 (1) |
| 4 | 0 (0) | 14 (6) | 14 (6) | 20 (9) | 4 (2) | 31 (13) | 15 (6) | 2 (2) |
| 5 | 19 (9) | 0 (1) | 0 (0) | 21 (10) | 46 (18) | 0 (0) | 0 (1) | 14 (6) |
| 6 | 5 (2) | 13 (5) | 21 (8) | 5 (3) | 3 (1) | 30 (12) | 19 (7) | 6 (3) |
| 7 | 2 (1) | 15 (5) | 16 (6) | 16 (6) | 4 (2) | 18 (7) | 24 (9) | 6 (2) |
| 8 | 9 (4) | 15 (6) | 1 (1) | 19 (7) | 18 (8) | 0 (0) | 5 (2) | 33 (13) |

**Fig. S4.**
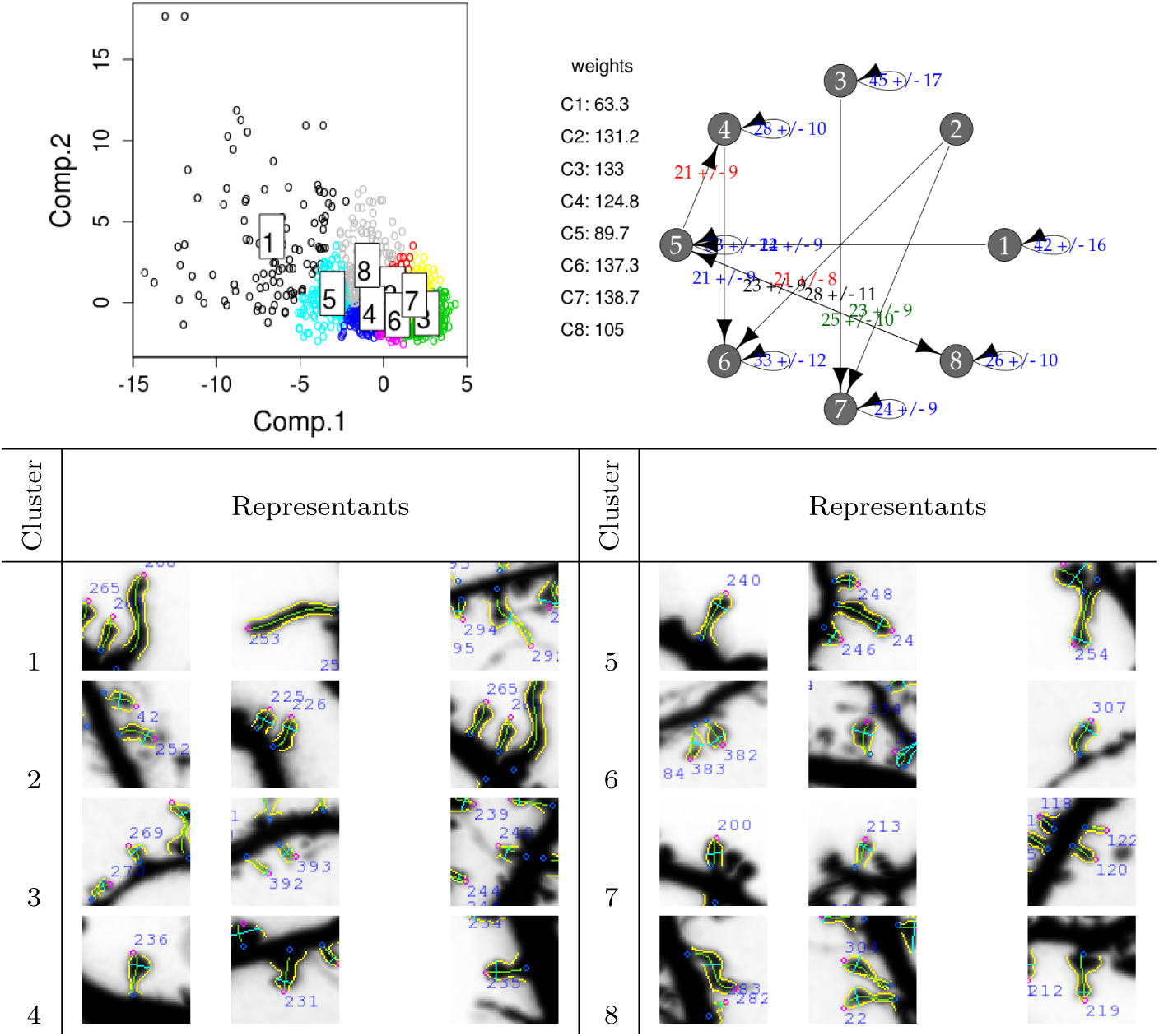
Results (clusters plot, *transition graph* and spines selected as clusters’ representants) of *cmeans* clustering applied to *ACTIVE* ∪ *CONTROL*. For each cluster the initial weight (sum of weights of spines in the cluster at time *t*_0_) is presented. Only transitions of values higher than 20% are shown on the graph. In contrast to *hierarchical* clustering case all clusters contain nonneglible amounts of spines. However, differences between spines from different clusters are not that significant and easy to interpret.

**Fig. S5.**
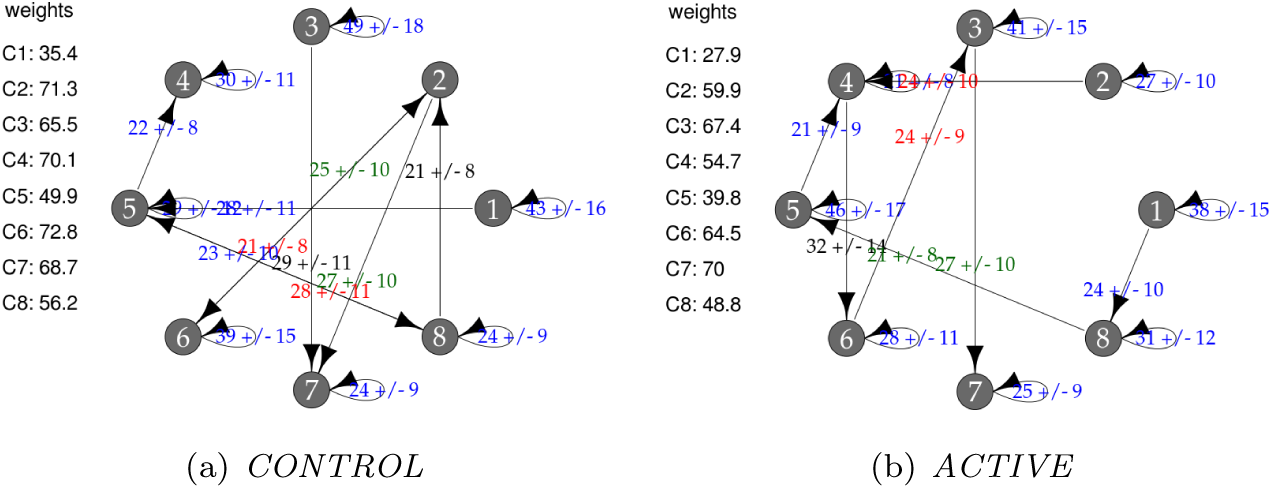
*Transition graphs* for for *cmeans* clustering. For each cluster the initial weight (sum of spines’ weights in the cluster at time *t*_0_) is presented. Values are given in rounded percents. Only transitions (probabilities) of values higher than 20% are shown. Subfigures should not be compared because they are computed for populations of different characteristic at *t*_0_.

**Fig. S6.**
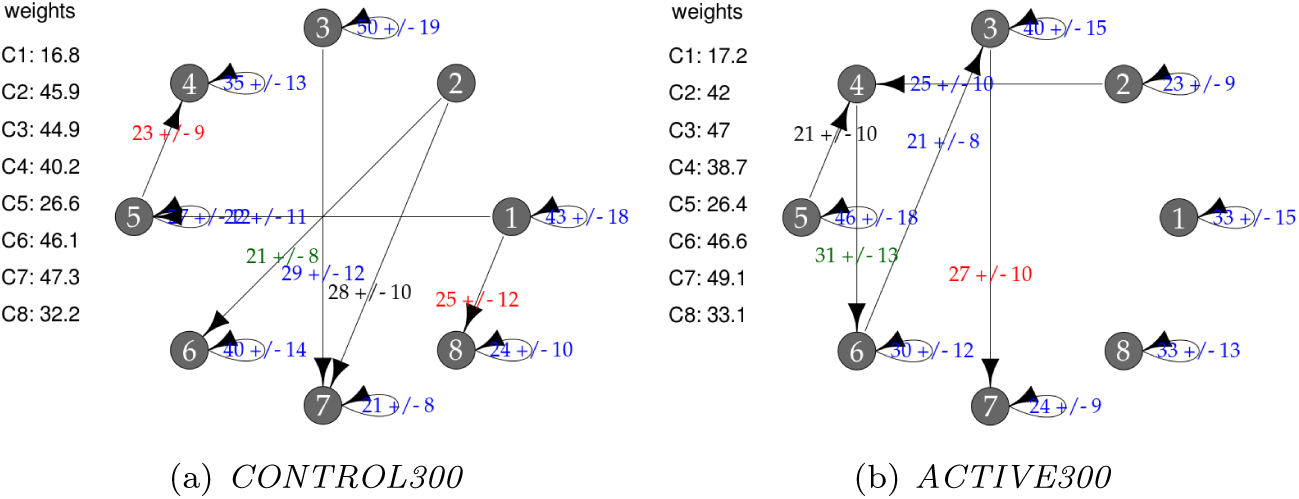
*Transition graphs* for balanced subpopulations and *cmeans* clustering. For each cluster the initial weight (sum of spines’ weights in the cluster at time *t*_0_) is presented. Values are given in rounded percents. Only transitions of values higher than 20% are shown. Although, most of observed differences between transitions are not significant when errors are taken into consideration, statistically significant difference between graphs was found using *RDC* statistic.

**Fig. S7.**
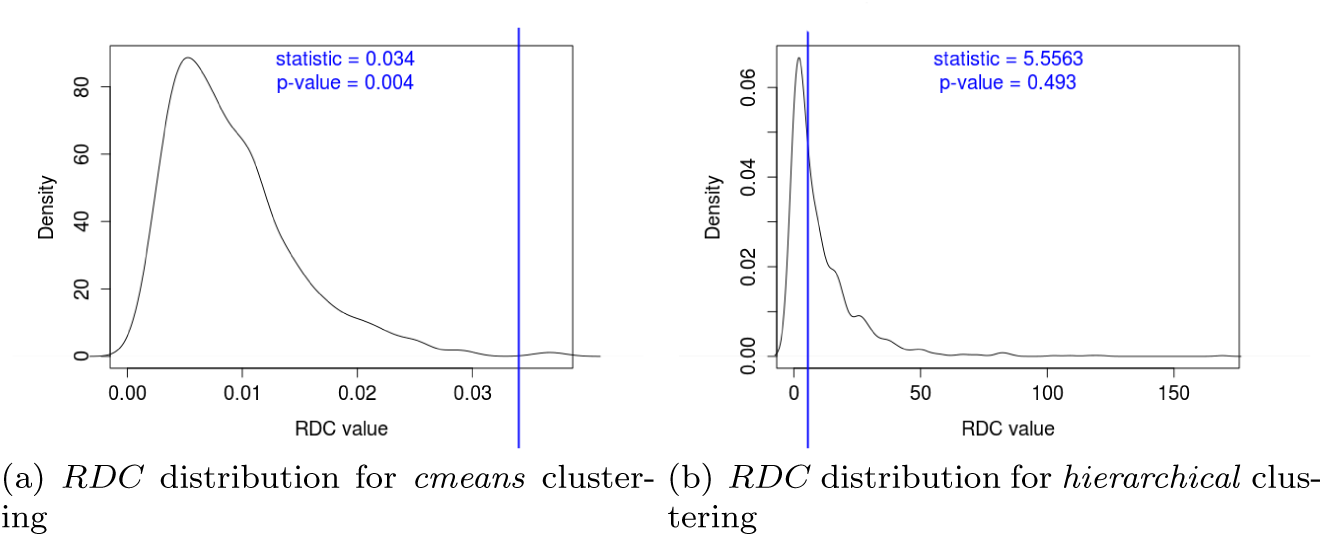
*RDC* statistic used to compare *CONTROL300* and *ACTIVE300*. Kernel estimation used for smoothing. Statistically significant difference between subpopulations observed for *cmeans* case.

**Fig. S8.**
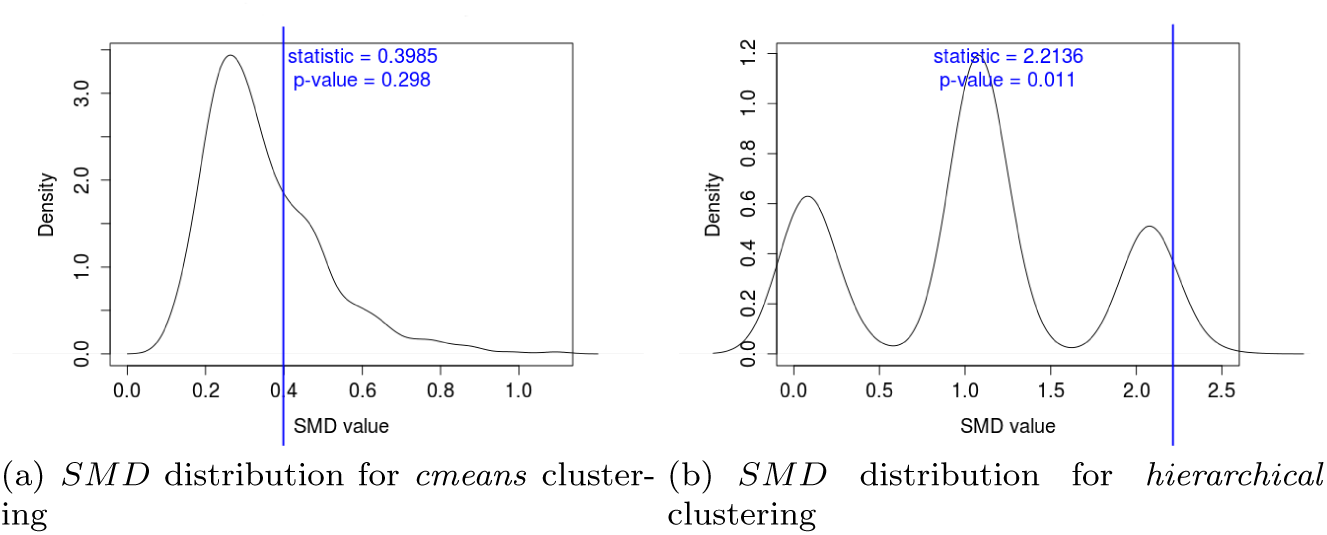
*SM D* statistic used to compare *CONTROL300* and *ACTIVE300*. Kernel estimation used for smoothing. Statistically significant difference between subpopulations observed for *hierarchical* case.

## Footnotes

1 https://bitbucket.org/3dome/spines

2 Even under ideal experimental conditions, variation among the spines is still present; thus, we choose to standardize them. This practice allows us to start with more homogeneous data and to reveal subtle differences.

3 The null hypothesis is that the means are equal, and the alternative is that the means are different.

4 Each feature is normalized by subtracting the mean and dividing by the standard deviation, both calculated based on the feature values from the sets.

5 We tried various different numbers of spines (100, 200, 300, 400) and concluded that 300 is the largest which satisfies the desired condition for spines closeness.

6 We expect that there is a higher percentage of growing spines from *ACTIVE* than from *CONTROL*. Therefore, we decide to define the threshold point between growing and not-growing groups to be the threshold maximizing the difference between number of growing spines from *ACTIVE* and the number of growing spines from *CONTROL*. What we observed for shrinking spines is similar, therefore we similarly seek the threshold point between shrinking and not-shrinking groups.

7 We expect that there is a higher percentage of growing spines from *ACTIVE* than from *CONTROL*. Therefore, we decide to define the threshold point between growing and not-growing groups to be the threshold maximizing the difference between number of growing spines from *ACTIVE* and the number of growing spines from *CONTROL*. What we observed for shrinking spines is similar, therefore we similarly seek the threshold point between shrinking and not-shrinking groups.

